# Robustness of Ancestral Recombination Graph Inference Tools to Phasing Errors

**DOI:** 10.1101/2025.11.24.690249

**Authors:** Leyan Wang, Yun Deng, Rasmus Nielsen

## Abstract

Ancestral Recombination Graphs (ARGs) are fundamental population genetic structures that encode the genealogical history of a sample of haplotypes along the genome. They have recently received substantial attention as they can be used to provide accurate estimates of population history, form the foundation for selection scans, provide time-labeled estimates of changes in mutation rates, and more. ARGs can be inferred using multiple different methods, but these methods all rely on computational haplotype phasing. Previous studies have shown that the accuracy of haplotype phasing can be low when done in small or medium-sized samples, which are typical for most non-model organisms. Conventional wisdom would, therefore, suggest that ARG inferences are of limited utility in most non-model organisms. To test this assumption, we benchmark the robustness of four ARG inference methods — Relate, SINGER, Threads, and tsinfer+tsdate — to phasing errors. Their performance with imperfect phasing was tested under simple and realistic demographic models, and evaluated in terms of estimated pairwise coalescence times, counts of inferred recombination breakpoints, and estimated branch-length-based diversity. We observe only a slight degradation in performance when using computational phasing across all four methods and all tested demographic scenarios, demonstrating a surprising robustness of ARG inference methods to phasing inaccuracies. These findings support broader application of ARG inference in settings where large reference panels are not available.

## 1 Introduction

The ancestral recombination graph (ARG) represents the genealogical history of a sample of haplotypes with recombination (Hudson, 1983; Griffiths and Marjoram, 1997). One simple way of thinking about ARGs is as the collection of coalescent trees in the entire genome. They can serve as a powerful tool for downstream coalescent-based population genetic analyses, including inference of demography (Speidel et al., 2019; Fan et al., 2025), selection (Stern et al., 2019, 2021; Hejase et al., 2022; Vaughn and Nielsen, 2024) and more. One of the reasons to focus on ARGs is that ARGs contain all the information in the data regarding demographic processes, and therefore, can form the basis for the development of full likelihood-based inference methods (Nielsen et al., 2025).

ARGs are not directly observed but must be inferred from genetic data. Accurate inference of ARGs is thus essential to our ability to fully exploit their potential. Recent advancements in ARG inference methods (Speidel et al., 2019; Kelleher et al., 2019; Gunnarsson et al., 2024; Deng et al., 2025) show promise in inferring high-quality ARGs for downstream applications, but these methods all require the availability of phased haplotypes. Genotype data produced using standard high-throughput sequencing or genotyping platforms is usually unphased, meaning that in two heterozygous sites, it is not determined which alleles are located on the same haplotype and which alleles are located on different haplotypes. The haplotypes cannot be directly inferred from the genotypes. However, the haplotypes can be estimated statistically when a sample of multiple individuals is available, or if a reference panel is available. This process is known as ‘haplotype phasing’. The accuracy of computational phasing methods has improved over the years, especially by taking advantage of large datasets and related individuals. However, computational phasing generally performs less well when only small samples of unrelated individuals are available (Browning and Browning, 2011; Marchini et al., 2006), as is often the case for non-model organisms. Phasing accuracy declines rapidly as sample size decreases and often requires thousands of individuals to reach optimal performance (Browning and Browning, 2007, 2011). Although the use of a reference panel is generally recommended to improve accuracy at small sample sizes (Loh et al., 2016), such panels are typically unavailable for non-model species. For cohorts with fewer than 100 individuals (200 haplotypes), the absence of a reference panel has been shown to substantially reduce phasing accuracy (Delaneau et al., 2013).

In this paper, we will evaluate the accuracy of ARG inference methods when haplotypes have been computationally phased in small samples. We will focus on two widely used computational phasing methods, Beagle (Browning et al., 2021) and SHAPEIT (Hofmeister et al., 2023), and explore how they perform in small samples and evaluate the downstream impact of phasing errors on ARG inferences. We focus on four ARG inference methods: Relate (Speidel et al., 2019), tsinfer (Kelleher et al., 2019) + tsdate (Wohns et al., 2022), SINGER (Deng et al., 2025), and Threads (Gunnarsson et al., 2024).

## 2 Methodology

We simulated ARGs and VCFs under a constant population size model, a CEU bottleneck model, and a selective sweep model. We artificially unphased the simulated VCFs and re-phased them with SHAPEIT and Beagle. We inferred ARGs using SINGER, Relate, Threads, and tsinfer+tsdate from both the simulated true VCFs and Beagle-phased VCFs and evaluated the quality of the ARG inference (Figure 1).

**Figure 1:**
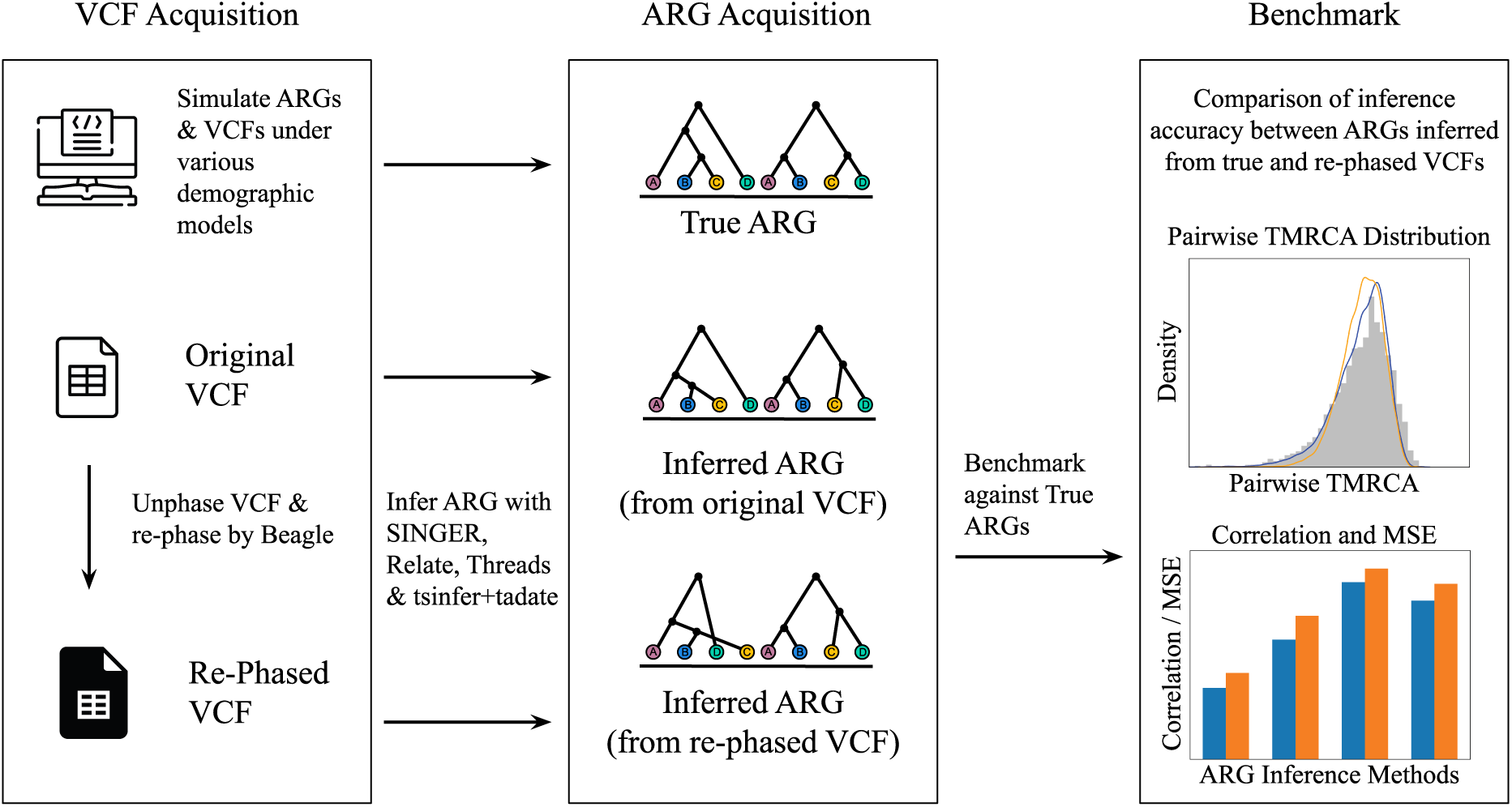
Workflow overview. Alt text: A schematic diagram showing a 3-step workflow: 1) VCF acquisition (original and re-phased VCFs), 2) ARG acquisition (from simulation, inferred from original VCFs, and inferrd from re-phased VCFs), and 3) benchmark inferred ARGs against true ARGs.

### 2.1 ARG and VCF Simulation

#### 2.1.1 Constant Population Size Model

We used msprime version 1.3.3 (Kelleher et al., 2016; Baumdicker et al., 2022) to simulate ARGs under the StandardCoalescent model (Hudson, 1983) with a constant effective population size of 10,000 diploid individuals, and a recombination rate of 2 *×* 10*^−^*^8^ per base pair per generation, for a total sequence length of 100 Mb. We generated VCFs of SNP data by simulating mutations on ARGs using a mutation rate of 2 *×* 10*^−^*^8^ per base pair per generation. Multiallelic sites were removed from the simulated VCFs.

#### 2.1.2 CEU Bottleneck Model

We simulated a sample of 50 diploid individuals with an inferred CEU demography (https://github.com/PalamaraLab/ASMC_data/tree/main/demographies) under the StandardCoalescent model with the other parameters being the same as the Constant Population Size Model.

#### 2.1.3 Selective Sweep Model

We performed simulations under the SweepGenicSelection model (Braverman et al., 1995; Kern and Schrider, 2016) followed by the StandardCoalescent model, using a sequence of length 10 Mb for 50 diploid individuals, with the other parameters being the same as the Constant Population Size Model. We set the beneficial allele in the middle of the genome at 5 Mb, with a selection coefficient of 0.02. The start frequency of the beneficial allele was 5 *×* 10*^−^*^6^ (= 1*/*2*N*_e_) and the end frequency was 0.5. We set the increment of time to 1 *×* 10*^−^*^6^. Sweep simulations in msprime proceed by first simulating a sweep trajectory, then simulating tree sequence and topology given the trajectory. Mutations are then subsequently placed on the trees according to a Poisson process. However, sequence data for the selected mutation are not directly simulated. Consequently, the msprime output does not label sequences with respect to carrier status of the selected mutation. To identify carrier haplotypes we re-simulate using the same seed for the random number generator, but stop the simulations at the time at which the beneficial allele arise. We then inspect the partial trees at the position of the beneficial allele, and identify the lineage which has approximately the same number of descendants in the full simulations as the reported derived allele frequency. While, in theory, this could leave some ambiguity regarding the identity of the carrier clade, we find that in all cases examined, the clade carrying the advantageous mutation is easily identified.

Once the carrier clade was identified, we manually inserted a mutation at the sweep position (5 Mb) into the simulated VCF. Haplotypes descending from the identified clade were assigned the derived allele, and all other haplotypes were assigned the ancestral allele.

### 2.2 Phasing Error Simulation

We first unphased the VCF files simulated by msprime by randomizing the order of alleles in each genotype. We then phased the unphased VCF files using Beagle (Browning et al., 2021) and SHAPEIT (Hofmeister et al., 2023).

#### 2.2.1 Beagle

We generated unphased genotype and Beagle-format map files from the unphased VCF files. We then ran Beagle version 5.5 with the map files to phase the haplotypes.

#### 2.2.2 SHAPEIT

We generated SHAPEIT-format map files from the unphased VCF files. Allele count (AC) and added allele number (AN) tags to the VCF files using bcftools +fill-tags from bcftools version 1.19 (Danecek et al., 2021). We then indexed the VCF files using bcftools index and ran phase common from SHAPEIT5 version 5.1.1 to phase the VCF files. Finally, we converted the output BCF files to VCF files using bcftools view.

#### 2.2.3 Phasing Error Rate

We used the switch error rate (SER) as a measure of phasing error rate, defined as the number of times two consecutive heterozygous sites were out of phase with each other over the total number of consecutive heterozygous sites. The phased VCFs were compared against the initial simulated VCFs and calculated SERs.

### 2.3 ARG Inference

We inferred ARGs from the simulated VCFs using four methods: Relate (Speidel et al., 2019, 2021), SINGER (Deng et al., 2025), Threads Gunnarsson et al., 2024, and tsinfer + tsdate (Kelleher et al., 2019; Wohns et al., 2022) using default parameters unless otherwise specified.

#### 2.3.1 Relate

We used Relate version 1.2.4 for ARG inference. We converted simulated VCF files to Relate haps and sample files using RelateFileFormats --mode ConvertFromVcf and inferred ARGs from haps and sample files with Relate --mode All, specifying the mutation rate (-m 2e-8), the haploid effective population size (-N 2e4), and a recombination map corresponding to a uniform recombination rate of 2 *×* 10*^−^*^8^ via --map. We converted Relate output to tskit Tree Sequence format using RelateFileFormats --mode ConvertToTreeSequence.

#### 2.3.2 SINGER

We ran SINGER beta version 0.1.8 using the command parallel singer, providing the diploid effective population size (-Ne 1e4) and the mutation rate (-m 2e-8). The default number of posterior samples (-n 100) and MCMC iterations between adjacent samples (-thin 20) were used.

#### 2.3.3 Threads

We converted simulated VCF files to the plink pgen and pvar formats using plink2 (Chang et al., 2015). We ran the command threads infer in Threads version 0.2.1, specifying -mutation rate 2e-8 and -recombination rate 2e-8. We specified a constant haploid effective population size of 20,000 using a demography file via --demography. Additionally, we provided a genetic map documenting uniform recombination along the genome via --map gz. Finally, we converted Threads output to tskit Tree Sequence format using threads convert.

#### 2.3.4 tsinfer and tsdate

We loaded simulated VCF files using cyvcf2.VCF and converted them to the tsinfer samples format, following the workflow described previously in the tsinfer tutorial (archived here: https://github.com/tskit-dev/tsinfer/commit/bb5355a). The tree sequences were inferred using tsinfer.infer from tsinfer version 0.5.1, specifying recombination rate = 2e-8. The genealogies inferred were simplified using TreeSequence.simplify from tskit version 0.5.8 (Kelleher et al., 2016; Wong et al., 2024) then dated by tsdate.date from tsdate version 0.2.1, specifying mutation rate = 2e-8.

#### 2.3.5 ARG Inference under Bottleneck Model

Tsinfer+tsdate does not accept need an effective population size as input and was run as described in the previous section. For ARG inference methods that require a single value of the effective population size as input (SINGER and Relate), we estimated an effective population size from the nucleotide diversity using *π* = 4 *· N*_e_ *· µ*. The Relate add-on module EstimatePopulationSize, which estimates effective population sizes and re-estimates ARG branch lengths, was applied to the Relate output. We note that running Relate without the add-on module also produced valid ARGs, but the distribution of inferred coalescence times was much less accurate, as they failed to capture the bimodality of the underlying distribution, despite showing similar correlation and mean squared errors to those inferred with the add-on module (see Supplementary Figure 5). Threads requires a demography file as an input. We ran Threads with a demography file describing a constant population size equal to the calculated value for a realistic benchmark, as detailed demographic history is rarely available in non-model organisms. However, we note that Threads also supports the use of a demography file with a detailed population size history. When run with detailed CEU demography, Threads produced ARGs with a more accurate distribution of inferred coalescence times that better captured the bimodality of the underlying distribution, with correlation and mean squared errors similar to those inferred under the constant population size demography (see supplementary Figure 5).

### 2.4 Benchmark

#### 2.4.1 Pairwise Coalescence Times Extraction for Distribution Visualization

We extracted pairwise coalescence times measured in number of generations at every 10kb along the genome, starting from the position of the first tree in the inferred ARGs. For each site along the genome, we sampled 20 pairs uniformly at random from all possible pairs, then extracted their pairwise coalescence times from true ARGs, ARGs inferred from original VCFs, and ARGs inferred from phased VCFs, using Tree.tmrca in tskit (Ralph et al., 2020). For SINGER, we selected a single posterior sample at random.

#### 2.4.2 Pairwise Coalescence Times Extraction for Correlation Calculation

We extracted pairwise coalescence times for all possible pairs of leaf nodes at 1,000 evenly spaced sites along the genome from true ARGs, ARGs inferred from original VCFs, and ARGs inferred from phased VCFs. For SINGER, we used an average over all posterior samples.

Comparison between true and inferred pairwise coalescence times requires that it is known, after phasing, which haplotype each allele is truly located on. However, when phasing the haplotypes using SHAPEIT or Beagle, this information is lost for homozygous sites. While each allele could, in theory, be labeled with its haplotype of origin, such labeling can, in practice, not be preserved when propagating the data through Beagle and SHAPEIT. We used two approaches to overcome this challenge. One approach was to only compare coalescence times in sites that were heterozygous in both individuals carrying the haplotypes, as the haplotype of origin is unambiguous for heterozygous sites. However, analyzing coalescence times in heterozygous sites only would lead to biased estimates. We therefore also devised a midpoint approach, described below, to reconstruct the local phase along the genome, allowing for comparison at any site along the genome. The midpoint approach may not fully recover the local phase, and comparisons based on this haplotype reconstruction method may produce mismatched pairs, resulting in overestimation of error rates. We, therefore, chose to present results from both approaches.

#### 2.4.3 Pairwise Coalescence Times at Heterozygous Sites

We extracted pairwise coalescence times from all pairs of individuals, where both individuals were heterozygous, for 1,000 randomly selected variable sites along the genome. For each pair of individuals, we extracted one haplotype from each individual and retrieved the pairwise coalescence time.

#### 2.4.4 Pairwise Coalescence Times with Local Phase Reconstruction

We devised a midpoint approach to reconstruct the local phase using information from heterozygous sites (Figure 2). At a given heterozygous site of a diploid individual, if the arrangement of alleles in the phased VCF was the same as the true phase in the simulated VCF, we recorded this pair of haplotypes as not switched at this position. If the arrangement was the opposite of the true phase, we recorded this pair of haplotypes as switched at this position. We compared the phased VCF with the original VCF to locate all switched heterozygous sites along the genome for each individual. Assuming there are no silent switches, we marked the region between two adjacent heterozygous sites as switched if both sites are switched and not switched if both sites are not switched. If the adjacent heterozygous sites have opposite switch status, we split the region at the midpoint between these two sites and marked each half to be the switch status of their closer heterozygous site. We marked the beginning and end of the genome as having the same switch status as the first and last heterozygous sites. We compared inferred pairwise coalescence times from re-phased VCFs to the true simulated coalescence times by matching re-phased haplotypes to the same pairs based on local switch states of the phased genome.

**Figure 2:**
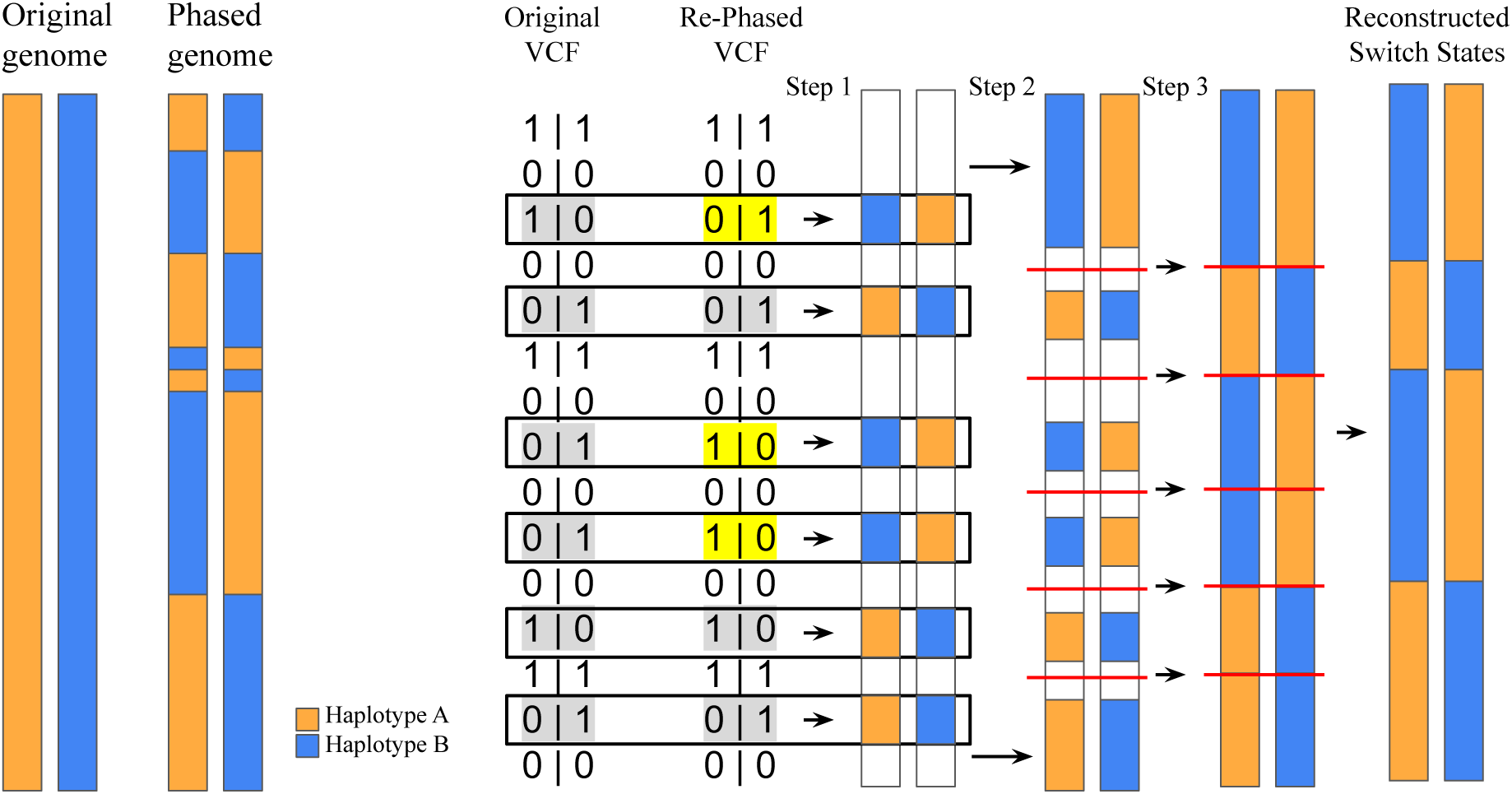
Methods for Local Phase Reconstruction. Step 1: Heterozygous sites are identified and compared between original and re-phased VCFs to determine whether the phase has switched at these positions. Step 2: The beginning and the end of the genome are marked as the same switch state as the first and the last heterozygous sites. Step 3: If the neighboring heterozygous sites are out of phase with each other, the switch state is split at the midpoint between these sites. All sites follow the same switch state as the closest heterozygous site. *Alt text: A schematic diagram showing the midpoint approach to local phase reconstruction*.

#### 2.4.5 Recombination Breakpoints

We obtained recombination breakpoints from simulated and inferred tree sequences using tskit TreeSequence.breakpoints (Ralph et al., 2020) and counted them. For SINGER, we used the average number of recombination points over all posterior samples.

#### 2.4.6 Branch-Length-Based Diversity Calculation

We calculated the branch-length-based diversity of carriers and non-carriers across 200 windows over the 1 Mb region around the site of selection using the branch mode of the TreeSequence.diversity method in tskit (Ralph et al., 2020). For SINGER, we used the average diversity over all posterior samples.

### 2.5 Empirical Analysis of *Anopheles gambiae*

For the empirical case study on *Anopheles gambiae*, we obtained the VCF from the Anopheles gambiae 1000 Genomes Consortium Ag1000G phase 1 AR3 data release (2015). Based on the iHS ranking on the Ag1000g Selection Atlas (gambiae 1000 Genomes Consortium, 2017b; Voight et al., 2006), we chose to analyze the cyp6p locus in the Uganda population which produced the highest iHS score. Within the cyp6p locus, we chose an arbitrary site among those showing a clear reduction in carrier diversity. Other sites produced similar results as documented in Supplementary Figure 7. We subsetted a VCF of Uganda individuals in the 1 Mb region around the cyp6p locus using bcftools (Danecek et al., 2021) and phased it by Beagle. We inferred ARGs from the phased VCF using SINGER with thin = 50. We estimated the selection coefficient and historic allele frequency trajectory using CLUES2 (Vaughn and Nielsen, 2024). Due to slow convergence, we burned in the first 20 samples when inferring with CLUES2. We performed all other downstream analyses using the tskit package.

## 3 Results

### 3.1 Accuracy of Computational Phasing Methods

We compared the accuracy of two computational phasing methods, SHAPEIT and Beagle, in a range of sample sizes from 20 to 200 haplotypes. Since SHAPEIT requires a minimum sample size of 100 haplotypes, we tested only for sample sizes above this threshold. At lower sample sizes, where only Beagle was tested, we observed that the switch error rate decreased rapidly as the sample size increased (Figure 3). At higher sample sizes, where both methods were tested, they produced indistinguishable switch error rates. Further increases in sample size yielded diminishing returns in terms of accuracy improvement for both methods.

**Figure 3:**
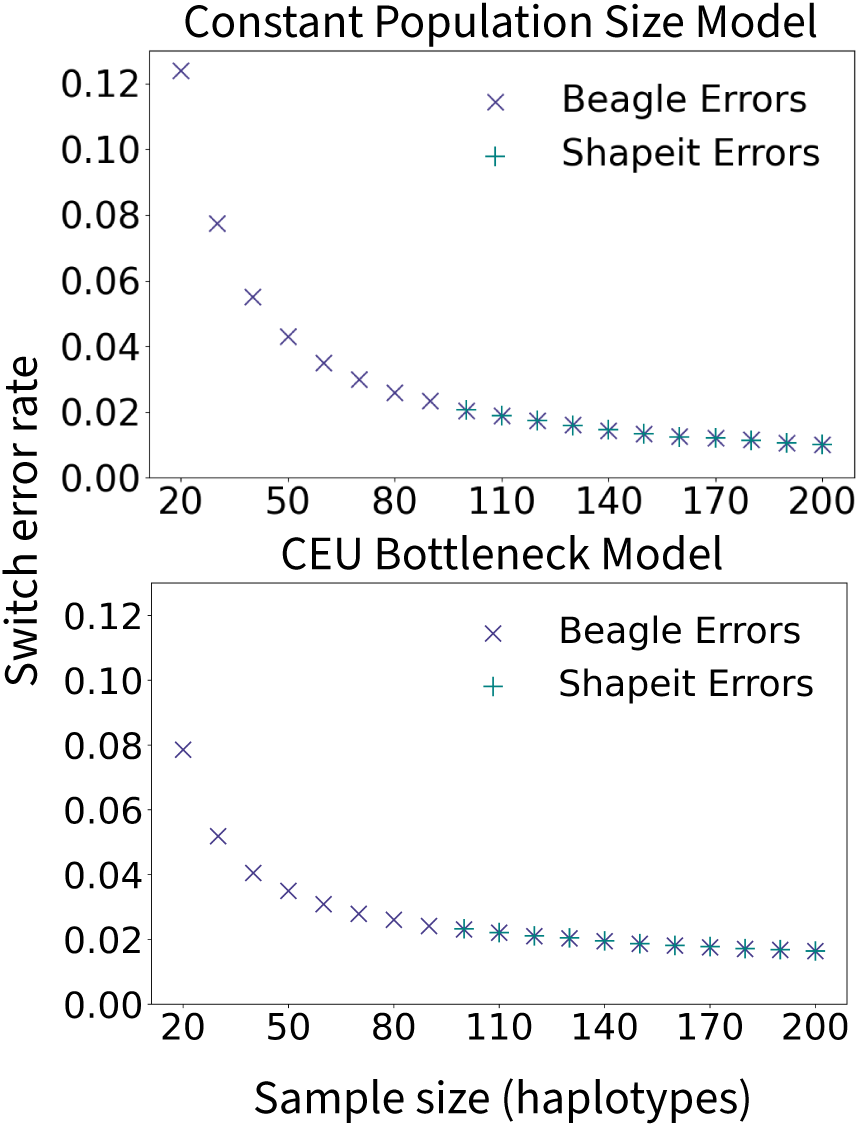
The switch error rate of statistical phasing tools over a range of sample sizes. The switch error rate was tested for simulations under the constant population size model (top) and the CEU bottleneck model (bottom). *Alt text: Scatter plots showing the switch error rate of Beagle and SHAPEIT over a range of sample sizes (10 to 200). The top plot shows error rates under a constant population size model and the bottom plot shows error rates under a CEU bottleneck model. Beagle and SHAPEIT showed indistinguishable performance and both showed decrease in error rates as sample sizes increase*.

### 3.2 Accuracy of ARG Inference with Computational Phasing

#### 3.2.1 Accuracy under Constant Population Size

ARGs and VCFs were simulated as described in Section 2 for sample sizes of 8, 20, and 100 haplotypes. Simulated VCFs were artificially unphased and then re-phased by Beagle, introducing phasing errors. ARGs were inferred from both true and Beagle-phased VCFs using four methods: SINGER, Relate, Threads, and tsinfer+tsdate. We compared the inferred coalescence times and recombination breakpoints to the simulated true values.

Pairwise coalescence times inferred from Beagle-phased VCFs were obtained based on haplotypes reconstructed using the midpoint approach as described in Section 2 and Figure 2. We compared the distribution of inferred pairwise coalescence times from true and Beagle-phased VCFs. At lower sample sizes, we observed underestimation of the most recent and the most ancient coalescence times and overestimation around the mode of the distribution (Figure 4.a). However, the difference became less pronounced as the sample size increased, and at a sample size of 100 haplotypes, the two distributions became indistinguishable, showing robustness of ARG inference methods to moderate phasing errors.

**Figure 4:**
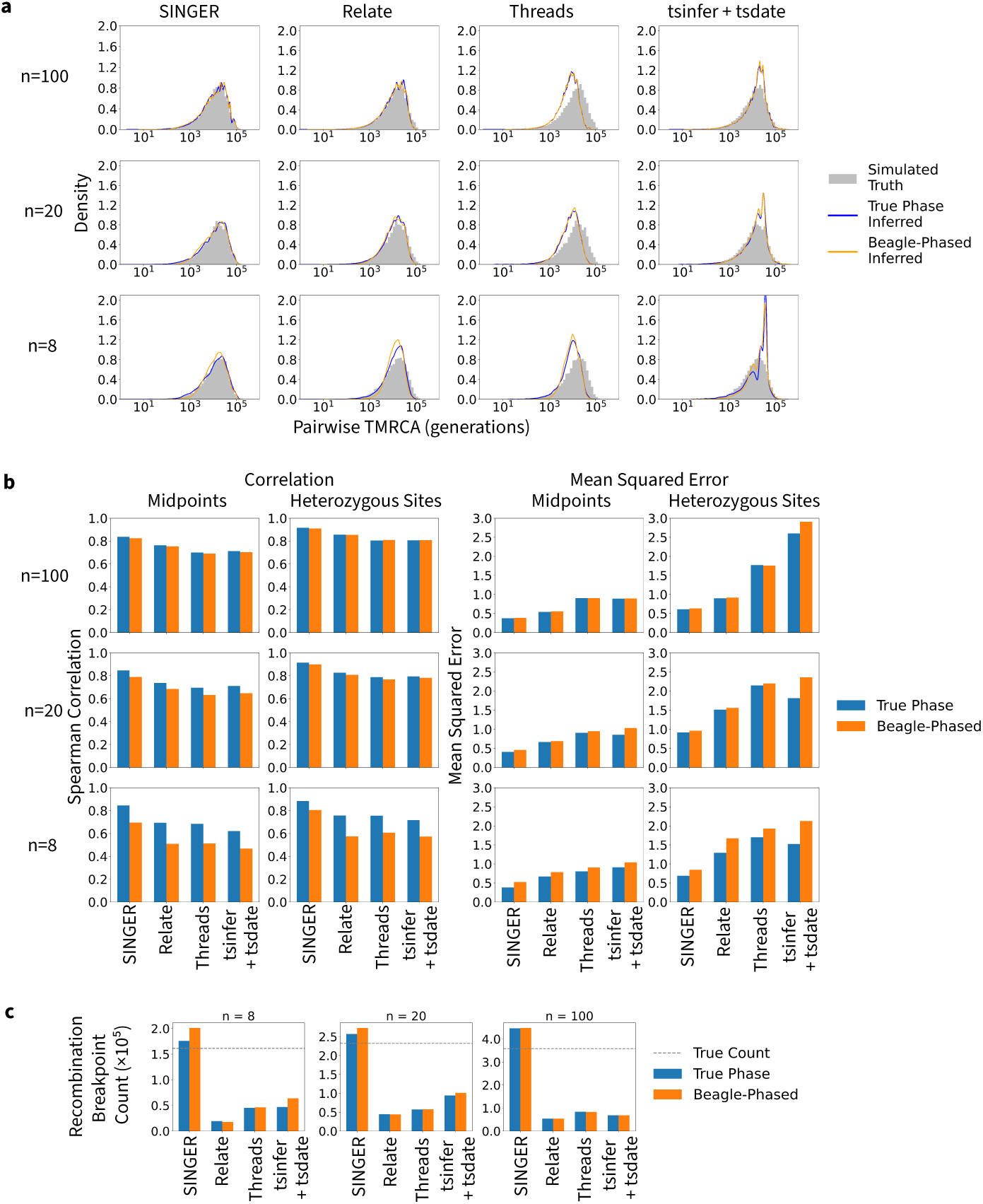
a. Distributions of pairwise coalescence times of true ARGs, ARGs inferred from true VCFs, and ARGs inferred from Beagle-Phased VCFs. b. The correlation (left half) and mean squared error (right half) between true and inferred ARGs based on midpoint switch state reconstruction and at heterozygous sites. c. The counts of inferred recombination breakpoints. *Alt text: A figure with panels labeled a to c. (a) A 3×4 grid of subpanels where each row shows*

We observed lower ARG inference quality after introduction of phasing errors as indicated by decreases in the Spearman correlation coefficient between true and inferred times and increases in mean squared error of inferred times (Figure 4.b). However, this gap in inference quality narrows *one sample size (100, 20, or 8) and each column shows an ARG inference method (SINGER, Relate, Threads, or tsinfer+tsdate). Each subpanel shows the coalescence times distributions for the simulation, ARGs inferred from true VCFs and ARGs inferred from Beagle-phased VCFs. (b) Two 3×2 grids of subpanels where each row shows one sample size (100, 20, or 8) and each column shows the comparison approach (midpoint or heterozygous sites). The two panels are bar plots showing the correlation and mean squared error respectively between simulations and ARGs inferred from true VCFs and Beagle-phased VCFs. (c) 3 bar plots showing the counts of recombination breakpoints at each sample size (8, 20, and 100) in simulated ARGs, ARGs inferred from true VCFs, and ARGs inferred from Beagle-phased VCFs.* as the sample size increases, becoming barely distinguishable at 100 haplotypes. We observed systematic trends of errors in inferred coalescence times for certain methods (Supplementary Figure 1 - 4). Threads systematically underestimates coalescence times for all sample sizes, while Tsinfer + tsdate systematically overestimates for all sample sizes. Most importantly, the difference in accuracy between methods is larger than the difference in accuracy using the true phasing and using computational phasing, as measured by the Spearman correlation coefficient or the mean squared error. Even for *n* = 8, the performance of Singer with computational phased haplotypes is better than that of Threads and tsinfer+tsdate with true haplotypes and comparable to that of Relate with true haplotypes.

Counts of inferred recombination breakpoints in the ARGs increased after the introduction of phasing errors (Figure 4.c; Supplementary Figure 6). Consistent with the trends observed for coalescence times inference, the difference diminished as the sample size increased and became mostly negligible for 100 haplotypes. More importantly, the effect of using phased haplotypes relative to true haplotypes is, in general, very minor compared to the biases already inherent in the inference methods. Relate, Threads and tsinfer+tsdate tends to strongly underestimate the number of recombination events. Singer tends to slightly overestimate the number of recombination events, possibly because the Bayesian estimator in this case is biased, or possibly because of mixing/convergence issues.

#### 3.2.2 Accuracy under CEU Bottleneck Demography

We simulated ARGs and VCFs under the CEU bottleneck demography and inferred ARGs from both true and Beagle-phased VCFs. The distributions of coalescence times inferred from true and Beagle-phased VCFs did not show much difference, consistent with observations under the constant population size at this sample size (Figure 5.a). The results differ very little in terms of Spearman correlation coefficient and MSE between ARGs inferred from true and Beagle-phased VCFs (Figure 5.b).

**Figure 5:**
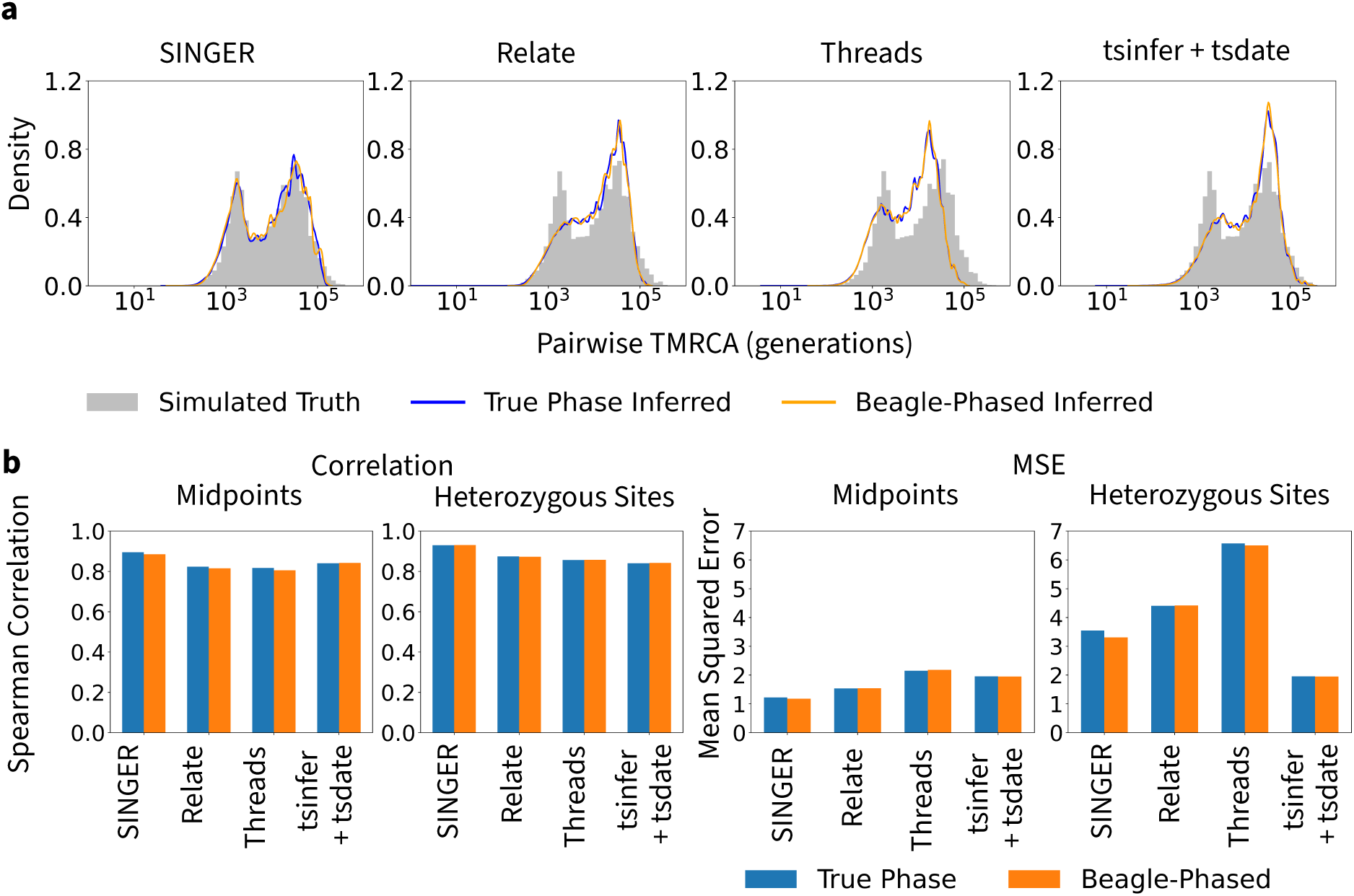
a. Distributions of pairwise coalescence times of true ARGs, ARGs inferred from true VCFs, and ARGs inferred from Beagle-Phased VCFs under the CEU bottleneck model. b. The correlation and mean squared error between true and inferred ARGs based on midpoint switch state reconstruction and at heterozygous sites. *Alt text: A figure with panels labeled a to b. (a) a panel with 4 subplots each showing the distribution of coalescence times of simulated ARGs, ARGs inferred from true VCFs, and ARGs inferred from Beagle-phased VCFs by each ARG inference method (SINGER, Relate, Threads and tsinfer+tsdate). (b) The left panel shows 2 bar plot of the correlation between true and inferred coalescence times for all 4 methods and for ARGs inferred from both true and Beagle-phased VCFs. One plot is calculated based on the reconstructed local phase (midpoint approach) and the other is calculated at heterozygous sites. The right panel shows 2 bar plot of the mean squared error between true and inferred coalescence times for all 4 methods and for ARGs inferred from both true and Beagle-phased VCFs. One plot is calculated based on the reconstructed local phase (midpoint approach) and the other is calculated at heterozygous sites*.

#### 3.2.3 Branch-Length-Based Diversity under the Selective Sweep Model

We simulated ARGs under the selective sweep model and calculated branch-length-based diversity of carriers and non-carriers of the beneficial allele along the genome. The carriers showed a clear reduction, relative to the non-carriers, in diversity around the site targeted by selection, as expected under a selective sweep.

The reduction of diversity around the selected site was well-preserved in all inferred ARGs (Figure 6). The magnitude of reduction was captured equally well in ARGs inferred from true VCFs and Beagle-phased VCFs, consistent with observations on pairwise coalescence times for this sample size (Figure 4). We note that ARGs inferred by Threads showed a systematic underestimation of branch-based diversity, consistent with previous results that showed systematic underestimation of coalescence times in Threads.

**Figure 6:**
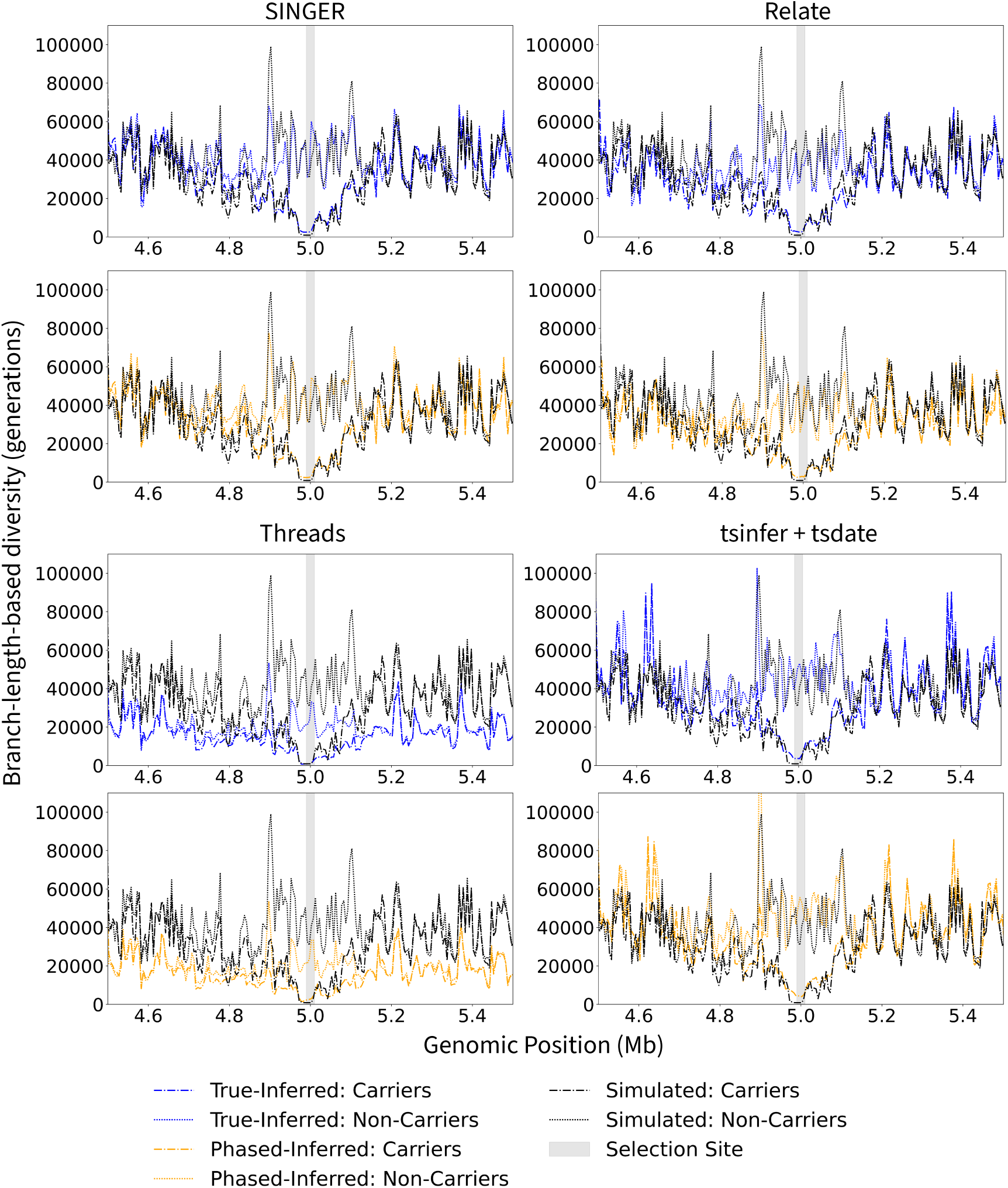
True and inferred branch-length-based diversity of carriers and non-carriers over the 1 Mb genomic region around the site of selection. For each method shown, the top plot shows ARG results inferred from true VCFs, the bottom plot shows ARG results inferred from Beagle-phased VCF. *Alt text: Line plots showing branch-length-based diversity among carriers vs non-carriers along a 1-megabase region around the mutation. For each method, there are one plot with diversity in ARGs inferred from true VCFs and one plot with diversity in ARGs inferred from Beagle-phased VCFs*.

### 3.3 ARGs Reveal Natural Selection Timeline at the *Anopheles gambiae* Cyp6p Locus

Natural selection in the *Anopheles gambiae* mosquitoes, vectors of the malaria parasite (*Plasmodium falciparum*), has been extensively studied due to concerns of pesticide resistance in malaria control (gambiae 1000 Genomes Consortium, 2017a; Davies et al., 2007; Clarkson et al., 2021; Mitchell et al., 2014; Edi et al., 2014). The cyp6p locus is one of the many loci that have been suspected to be under strong positive selection due to pesticide usage (Edi et al., 2014). However, previous summary statistics methods did not identify the time course of selection, and it is still unknown if the selection in cyp6p locus can be directly connected to recent selection in response to pesticide use. Studies comparing samples collected at different times in the last few decades (Clarkson et al., 2021; Kengne-Ouafo et al., 2026), tend to find mutations already segregating at high frequencies, suggesting selection acting on standing variation, and perhaps selection starting before the use of pesticides. It remains difficult to trace the origin and allele frequency trajectories of mutations of interests. To address this question, we infer local ARGs with SINGER (Deng et al., 2025), and reconstruct the historical allele frequency with CLUES2 (Stern et al., 2019; Vaughn and Nielsen, 2024), revealing the temporal dynamics of natural selection on the cyp6p locus.

Analysis with inferred ARGs recovered a significant signal of strong positive selection, consistent with previous evidence of selection at this locus (Figure 7.a). We calculated the tree-based statistic for detecting positive selection proposed by Speidel et al. (2019) with the local tree in the last posterior sample from SINGER, which suggests a highly significant signal of natural selection (p-value = 2.52 *×* 10*^−^*^4^). Similarly, CLUES2 estimated significant strong positive selection with a selection coefficient of 0.00819 over the last 200,000 generations (p-value = 5.75 *×* 10*^−^*^171^) from 80 posterior samples. Visual patterns in the ARG provided complementary evidence. The branch-length-based diversity of carriers of the derived variant showed a notable decrease around the site compared to non-carriers (Figure 7.a). The inferred local tree also showed rapid expansion of the carrier lineage over time compared to other neutral lineages (Figure 7.c). We note that the causal SNP in cyp6p is not known, and we therefore investigated robustness of our results to the choice of SNP. As shown in Supplementary Figure 7, other potential candidate SNPs, than the one chosen in Figure 7, show similar patterns.

**Figure 7:**
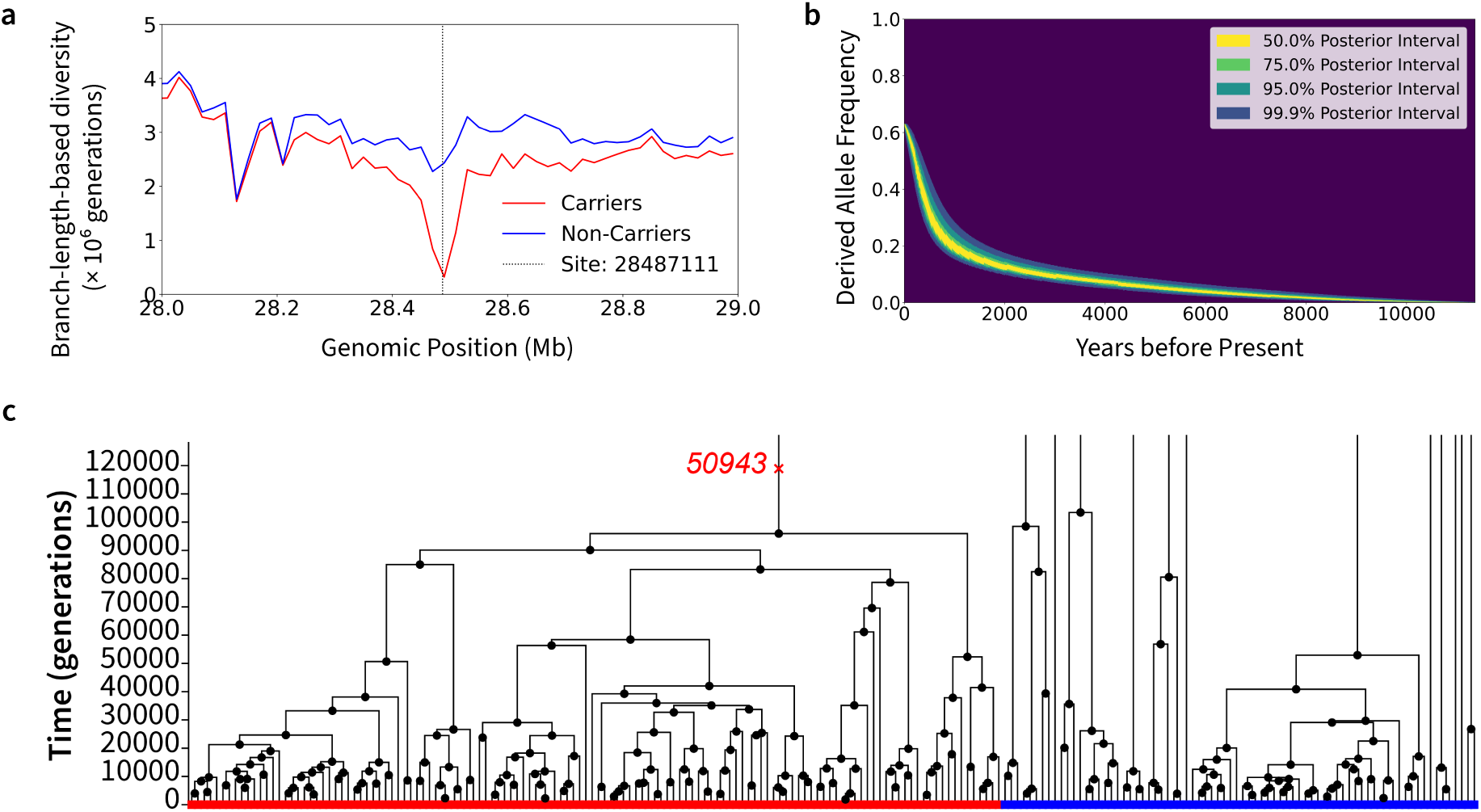
a. The branch-length-based diversity of carriers and non-carriers over the 1 Mb genomic region around the site of selection in cyp6p. b. The CLUES2-estimated allele frequency trajectory of the mutation under selection. c. The local tree at the site of selection. *Alt text: A figure with 3 panels a to c. (a) A line graph of the branch-length-based diversity of carriers and non-carriers of the mutation at site 28,487,111 along the 1-megabase region surrounding it. (b) A shaded confidence band plot showing the 50%, 75%, 95%, and 99.9% posterior interval of the CLUES2-estimated drived allele frequency over the last 11,000 years. (c) A diagram showing the local tree topology at site 28,487,111 in the past 120,000 generations*.

Contrary to previous beliefs that the natural selection in African mosquito population is caused by recent pesticide usage, the ARG results suggested selection that started well before the use of pesticide. The mutation we studied appears to be a standing variant. Using 80 posterior samples from SINGER, we dated the age of the mutation to around 13,656 years ago (95% posterior predictive interval = [8710, 178291], Supplementary Figure 8). The mutation age is consistent with the observation of an estimated average coalescence time of *π/*(2*µ*) = 8, 699 years, when calculated among carriers in a 20kb region around the mutation. Here *π* and (*µ*) is the nucleotide diversity and mutation rate, respectively. The allele frequency trajectory estimated by CLUES2 showed an increase in frequency starting around a thousand years ago (Figure 7.b). An ancient onset is also supported by the shallow and narrow shape of the trough in the diversity curve (Figure 7.a). More recent selection would have produced a wider and deeper trough.

## 4 Discussion

We evaluated common methods for ARG inference when haplotypes have been obtained using computational phasing. We generally found that the reduction in accuracy in terms of inference of coalescence times, or recombination rates, is small when haplotypes have been computationally phased. The difference between computational methods for inference of ARGs is much larger than the difference in performance when applying the methods to true haplotypes versus computationally inferred haplotypes. This is also true even for surprisingly small samples of 8 haplotypes. The reason is likely that phasing for common alleles can be done very well even in small samples, and these alleles are most important for the inference of the genealogy. Rare mutations that are harder to phase may affect ARG inferences less.

Hitherto, most applications of ARG inferences have been in humans. We surmise that this is because of a hesitancy to apply ARG inference methods to cases where large reference samples for haplotype inference are not available. Our results suggest that the field should be encouraged to apply ARGs even to systems with limited phasing quality. Our case study on Uganda malaria mosquitoes (*Anopheles gambiae*) exemplifies the promise of applying ARGs to such systems. With ARGs inferred from reference-free computational phasing data, we revealed that a selection signal thought to be driven by recent pesticide usage, in fact started before the onset of pesticide usage. This illustrates the usefulness of ARG to provide temporal analyses, bringing potential for new discoveries of past events.

While ARG inference shows strong robustness to phasing errors, they may not necessarily perform as well in the presence of other sources of errors. Population genetic inferences are challenging due to assembly errors in reference genomes, ambiguity in alignment of short reads, contamination, sequencing errors, and more. They also typically rely on many assumptions regarding population structure and history, mutation and recombination processes, absence/presence of natural selection, breeding structure, and introgression with other species. While further investigation is required to fully understand the extend to which ARG inference may be affected by other uncertainties, our results demonstrate that the usual level of computational phasing errors does not compromise the performance of the ARG inference methods tested. Therefore, we encourage the field to apply ARG inference methods to systems previously shadowed by concerns over small sample sizes and limited phasing quality.

## 5 Data Availability

The code and data used in this article are available on GitHub at https://github.com/wleyan/ ARG-and-phasing.

## 6 Acknowledgment

We thank Yan Wong for helpful guidance on running tsinfer. This research was supported by NIH grant R35GM153400.

## References

Baumdicker, F., Bisschop, G., Goldstein, D., Gower, G., Ragsdale, A. P., Tsambos, G., Zhu, S., Eldon, B., Ellerman, E. C., Galloway, J. G., Gladstein, A. L., Gorjanc, G., Guo, B., Jeffery, B., Kretzschumar, W. W., Lohse, K., Matschiner, M., Nelson, D., Pope, N. S., Quinto-Cortés, C. D., Rodrigues, M. F., Saunack, K., Sellinger, T., Thornton, K., van Kemenade, H., Wohns, A. W., Wong, Y., Gravel, S., Kern, A. D., Koskela, J., Ralph, P. L., and Kelleher, J. (2022). Efficient ancestry and mutation simulation with msprime 1.0. Genetics, 220(3):iyab229.

Braverman, J. M., Hudson, R. R., Kaplan, N. L., Langley, C. H., and Stephan, W. (1995). The hitchhiking effect on the site frequency spectrum of dna polymorphisms. Genetics, 140(2):783.

Browning, B. L., Tian, X., Zhou, Y., and Browning, S. R. (2021). Fast two-stage phasing of large-scale sequence data. The American Journal of Human Genetics, 108(10):1880–1890.

Browning, S. R. and Browning, B. L. (2007). Rapid and accurate haplotype phasing and missing-data inference for whole-genome association studies by use of localized haplotype clustering. The American Journal of Human Genetics, 81(5):1084–1097.

Browning, S. R. and Browning, B. L. (2011). Haplotype phasing: existing methods and new developments. Nature Reviews Genetics, 12(10):703–714.

Chang, C. C., Chow, C. C., Tellier, L. C., Vattikuti, S., Purcell, S. M., and Lee, J. J. (2015). Second-generation plink: Rising to the challenge of larger and richer datasets. GigaScience, 4(1):s13742–015–0047–8.

Clarkson, C. S., Miles, A., Harding, N. J., O’Reilly, A. O., Weetman, D., Kwiatkowski, D., Donnelly, M. J., and gambiae 1000 Genomes Consortium, A. (2021). The genetic architecture of target-site resistance to pyrethroid insecticides in the african malaria vectors anopheles gambiae and anopheles coluzzii. Molecular ecology, 30(21):5303–5317.

Danecek, P., Bonfield, J. K., Liddle, J., Marshall, J., Ohan, V., Pollard, M. O., Whitwham, A., Keane, T., McCarthy, S. A., Davies, R. M., and Li, H. (2021). Twelve years of samtools and bcftools. GigaScience, 10(2):giab008.

Davies, T., Field, L., Usherwood, P., and Williamson, M. (2007). A comparative study of voltage-gated sodium channels in the insecta: implications for pyrethroid resistance in anopheline and other neopteran species. Insect molecular biology, 16(3):361–375.

Delaneau, O., Zagury, J.-F., and Marchini, J. (2013). Improved whole-chromosome phasing for disease and population genetic studies. Nature methods, 10(1):5–6.

Deng, Y., Nielsen, R., and Song, Y. S. (2025). Robust and accurate bayesian inference of genome-wide genealogies for hundreds of genomes. Nature genetics, 57(9):2124–2135.

Edi, C. V., Djogbenou, L., Jenkins, A. M., Regna, K., Muskavitch, M. A., Poupardin, R., Jones, C. M., Essandoh, J., Ketoh, G. K., Paine, M. J., et al. (2014). Cyp6 p450 enzymes and ace-1 duplication produce extreme and multiple insecticide resistance in the malaria mosquito anopheles gambiae. PLoS genetics, 10(3):e1004236.

Fan, C., Cahoon, J. L., Dinh, B. L., Ortega-Del Vecchyo, D., Huber, C. D., Edge, M. D., Mancuso, N., and Chiang, C. W. (2025). A likelihood-based framework for demographic inference from genealogical trees. Nature genetics, 57(4):865—-874.

gambiae 1000 Genomes Consortium, A. (2017a). Genetic diversity of the african malaria vector anopheles gambiae. Nature, 552(7683):96.

gambiae 1000 Genomes Consortium, T. A. (2015). Ag1000g phase 1 ar3 data release.

gambiae 1000 Genomes Consortium, T. A. (2017b). Ag1000g selection atlas. Accessed: 2026-07-29.

Griffiths, R. C. and Marjoram, P. (1997). An ancestral recombination graph. Institute for Mathematics and its Applications, 87:257.

Gunnarsson, Á. F., Zhu, J., Zhang, B. C., Tsangalidou, Z., Allmont, A., and Palamara, P. F. (2024). A scalable approach for genome-wide inference of ancestral recombination graphs. bioRxiv. 2024.08.31.610248.

Hejase, H. A., Mo, Z., Campagna, L., and Siepel, A. (2022). A deep-learning approach for inference of selective sweeps from the ancestral recombination graph. Molecular Biology and Evolution, 39(1):msab332.

Hofmeister, R. J., Ribeiro, D. M., Rubinacci, S., and Delaneau, O. (2023). Accurate rare variant phasing of whole-genome and whole-exome sequencing data in the UK Biobank. Nature Genetics, 55(7):1243–1249.

Hudson, R. R. (1983). Properties of a neutral allele model with intragenic recombination. Theoretical population biology, 23(2):183–201.

Kelleher, J., Etheridge, A. M., and McVean, G. (2016). Efficient coalescent simulation and genealogical analysis for large sample sizes. PLoS computational biology, 12(5):e1004842.

Kelleher, J., Wong, Y., Wohns, A. W., Fadil, C., Albers, P. K., and McVean, G. (2019). Inferring whole-genome histories in large population datasets. Nature Genetics, 51(9):1330–1338.

Kengne-Ouafo, J. A., Kouamo, M. F., Muhammad, A., Tepa, A., Ntadoun, S., Mugenzi, L. M., Tekoh, T., Hearn, J., Tchouakui, M., Wondji, M., et al. (2026). The e205d mutation in cyp6p3 drives pyrethroid insecticide resistance in the african malaria mosquito vector anopheles gambiae. Science Translational Medicine, 18(835):eado6222.

Kern, A. D. and Schrider, D. R. (2016). Discoal: Flexible coalescent simulations with selection. Bioinformatics, 32(24):3839–3841.

Loh, P.-R., Danecek, P., Palamara, P. F., Fuchsberger, C., A Reshef, Y., K Finucane, H., Schoenherr, S., Forer, L., McCarthy, S., Abecasis, G. R., et al. (2016). Reference-based phasing using the haplotype reference consortium panel. Nature genetics, 48(11):1443–1448.

Marchini, J., Cutler, D., Patterson, N., Stephens, M., Eskin, E., Halperin, E., Lin, S., Qin, Z. S., Munro, H. M., Abecasis, G. R., et al. (2006). A comparison of phasing algorithms for trios and unrelated individuals. The American Journal of Human Genetics, 78(3):437–450.

Mitchell, S. N., Rigden, D. J., Dowd, A. J., Lu, F., Wilding, C. S., Weetman, D., Dadzie, S., Jenkins, A. M., Regna, K., Boko, P., et al. (2014). Metabolic and target-site mechanisms combine to confer strong ddt resistance in anopheles gambiae. PloS one, 9(3):e92662.

Nielsen, R., Vaughn, A. H., and Deng, Y. (2025). Inference and applications of ancestral recombination graphs. Nature Reviews Genetics, 26(1):47–58.

Ralph, P., Thornton, K., and Kelleher, J. (2020). Efficiently summarizing relationships in large samples: A general duality between statistics of genealogies and genomes. Genetics, 215(3):779–797.

Speidel, L., Cassidy, L., Davies, R. W., Hellenthal, G., Skoglund, P., and Myers, S. R. (2021). Inferring population histories for ancient genomes using genome-wide genealogies. Molecular Biology and Evolution, 38(9):3497–3511.

Speidel, L., Forest, M., Shi, S., and Myers, S. R. (2019). A method for genome-wide genealogy estimation for thousands of samples. Nature Genetics, 51(9):1321–1329.

Stern, A. J., Speidel, L., Zaitlen, N. A., and Nielsen, R. (2021). Disentangling selection on geneti-cally correlated polygenic traits via whole-genome genealogies. The American Journal of Human Genetics, 108(2):219–239.

Stern, A. J., Wilton, P. R., and Nielsen, R. (2019). An approximate full-likelihood method for inferring selection and allele frequency trajectories from dna sequence data. PLoS Genetics, 15(9):e1008384.

Vaughn, A. H. and Nielsen, R. (2024). Fast and accurate estimation of selection coefficients and allele histories from ancient and modern dna. Molecular biology and evolution, 41(8):msae156.

Voight, B. F., Kudaravalli, S., Wen, X., and Pritchard, J. K. (2006). A map of recent positive selection in the human genome. PLoS biology, 4(3):e72.

Wohns, A. W., Wong, Y., Jeffery, B., Akbari, A., Mallick, S., Pinhasi, R., Patterson, N., Reich, D., Kelleher, J., and McVean, G. (2022). A unified genealogy of modern and ancient genomes. Science, 375(6583). eabi8264.

Wong, Y., Ignatieva, A., Koskela, J., Gorjanc, G., Wohns, A. W., and Kelleher, J. (2024). A general and efficient representation of ancestral recombination graphs. Genetics, 228(1):iyae100.

