## Supplementary Materials for "Robustness of Ancestral Recombination Graph Inference Tools to Phasing Errors"

### 1 Heatmaps of Inferred vs True Coalescence Times

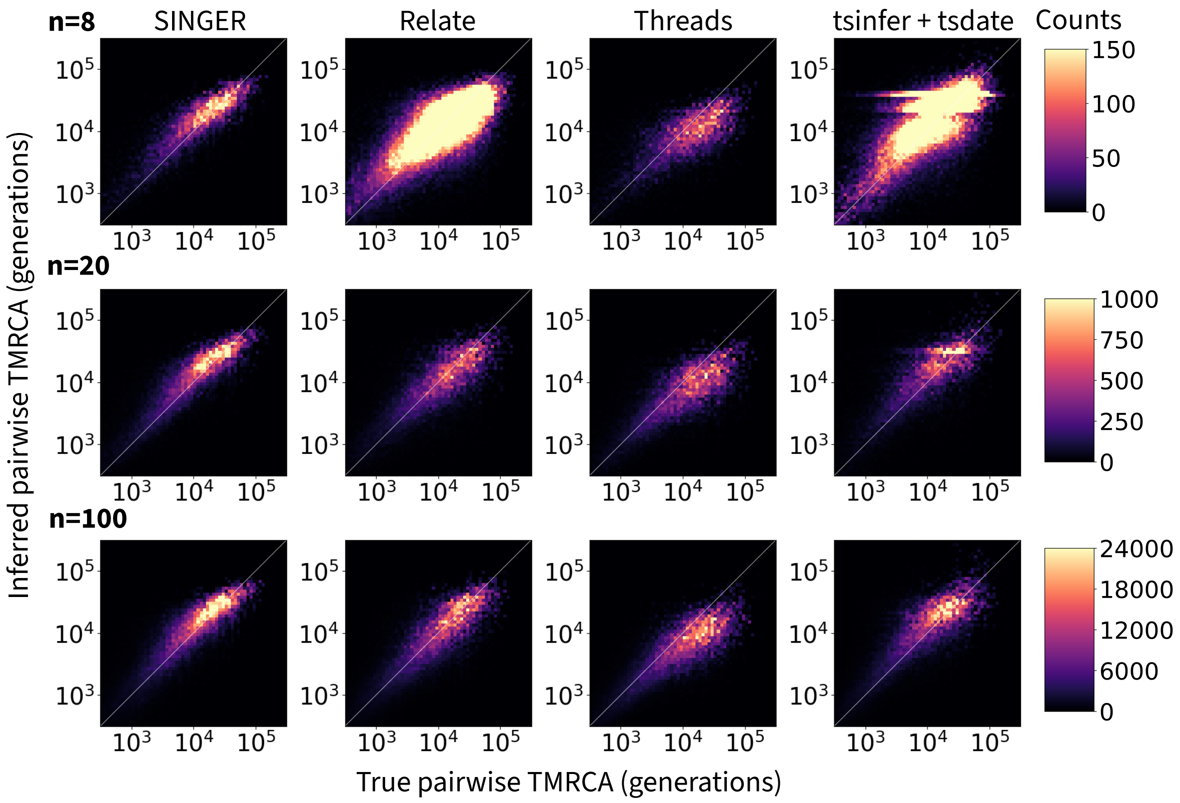


Figure 1: Pairwise coalescence times inferred from true VCFs compared with simulated ground truths for all sample sizes under the constant population size model.

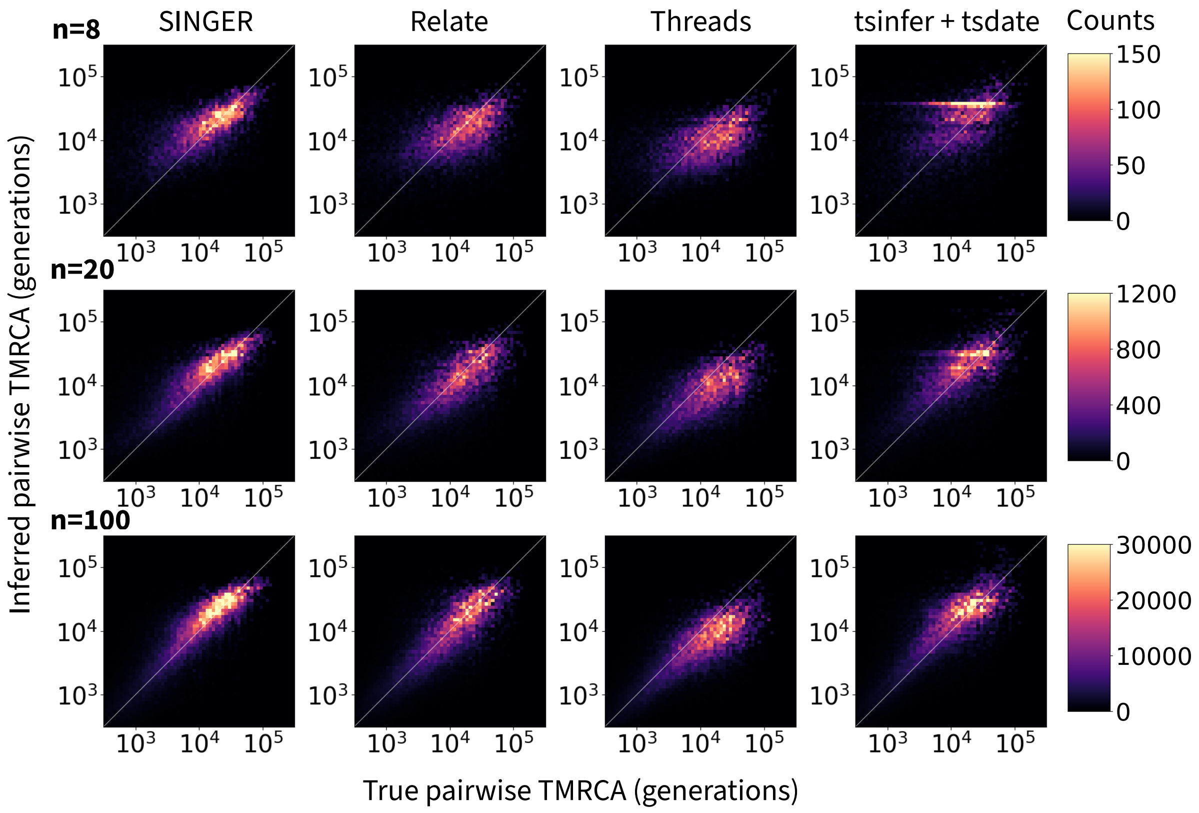


Figure 2: Pairwise coalescence times inferred from Beagle-Phased VCFs and adjusted using the midpoint approach compared with simulated ground truths for all sample sizes under the constant population size model.

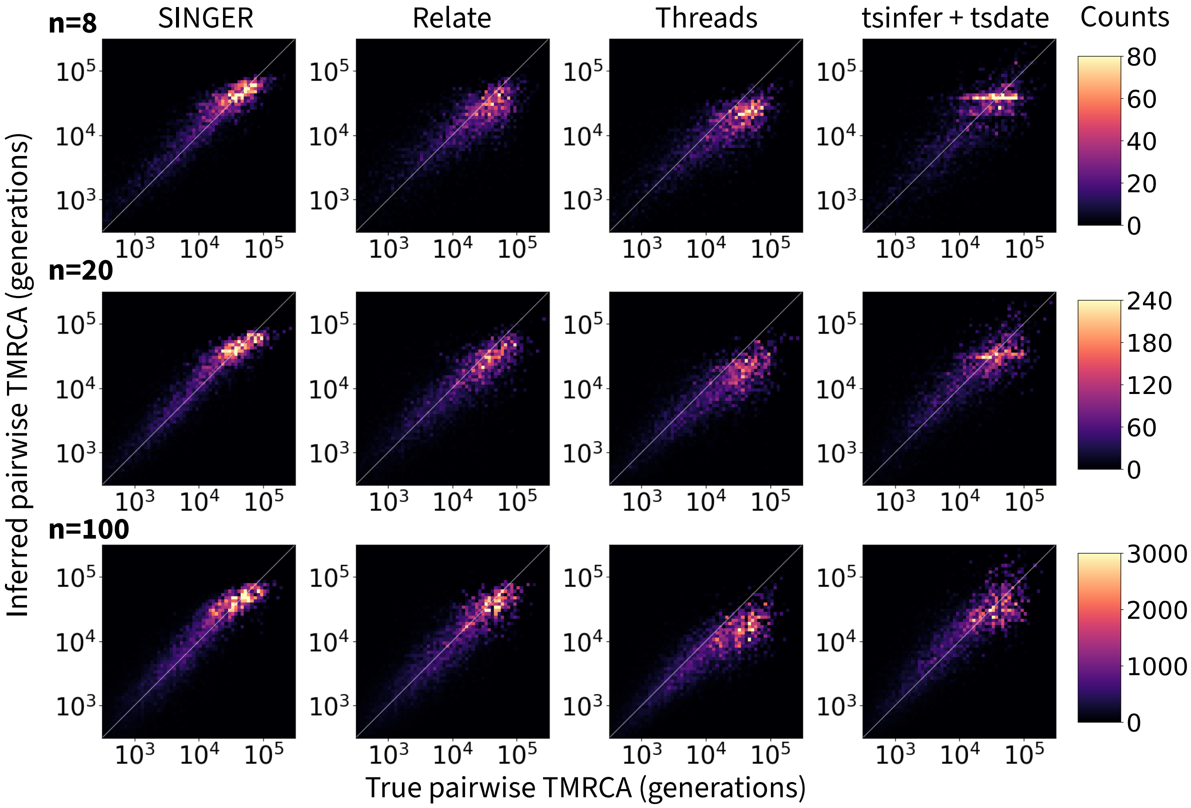


Figure 3: Pairwise coalescence times at heterozygous sites inferred from true VCFs compared with simulated ground truths for all sample sizes under the constant population size model.

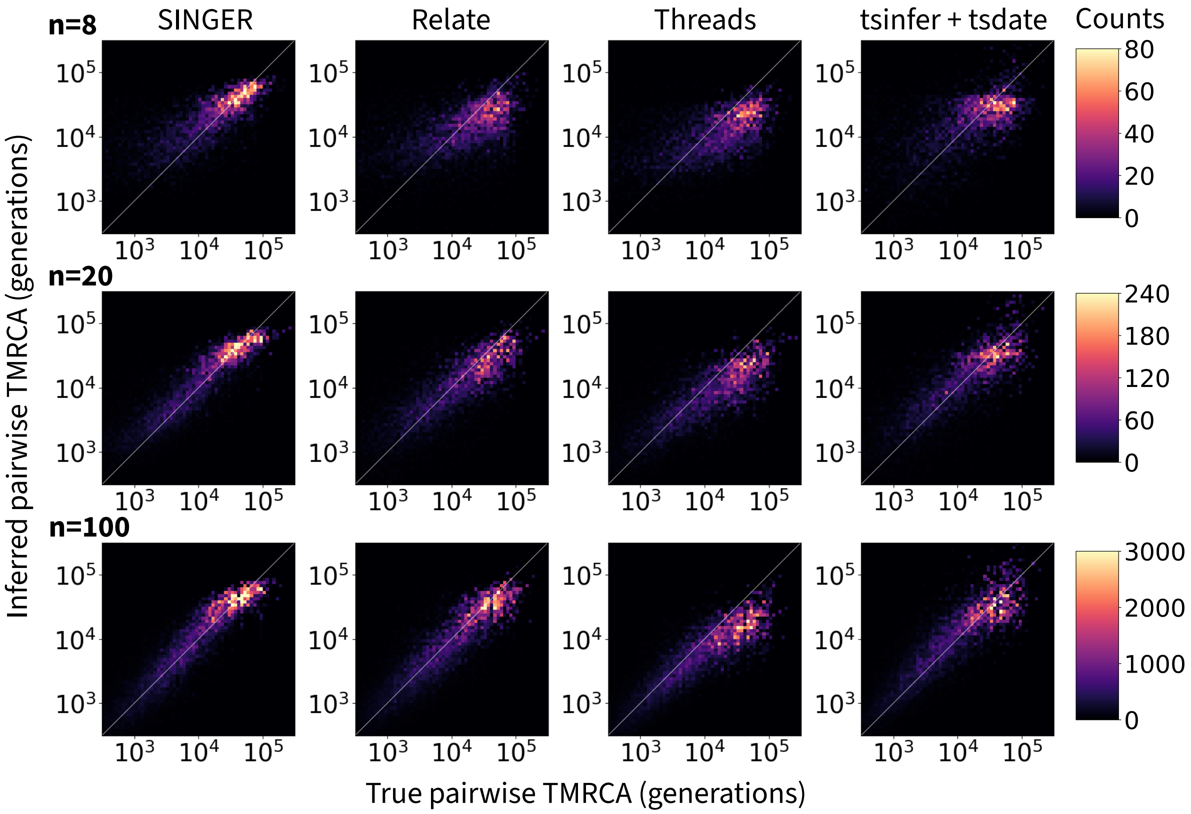


Figure 4: Pairwise coalescence times at heterozygous sites inferred from Beagle-phased VCFs compared with simulated ground truths for all sample sizes under the constant population size model.

### 2 Alternative CEU Model Specification for Relate and Threads

Relate offers an add-on module `EstimatePopulationSize` which estimates effective population sizes and re-estimates branch lengths to produce an updated ARG. For ARG inference under the CEU bottleneck model, we compared ARGs inferred with and without this add-on module. We confirmed that applying the `EstimatePopulationSize` module improved ARG inference accuracy and recovered the bimodality of the distribution of pairwise coalescence times better. Threads offers the option to supply a detailed demography file during inference. We decided it would be unrealistic for most researchers to provide such detailed demographic history, and thus supplied Threads with only a constant population size in the main benchmark. Here, we present the theoretical best performance when Threads was supplied with the same demography file used in the simulation.

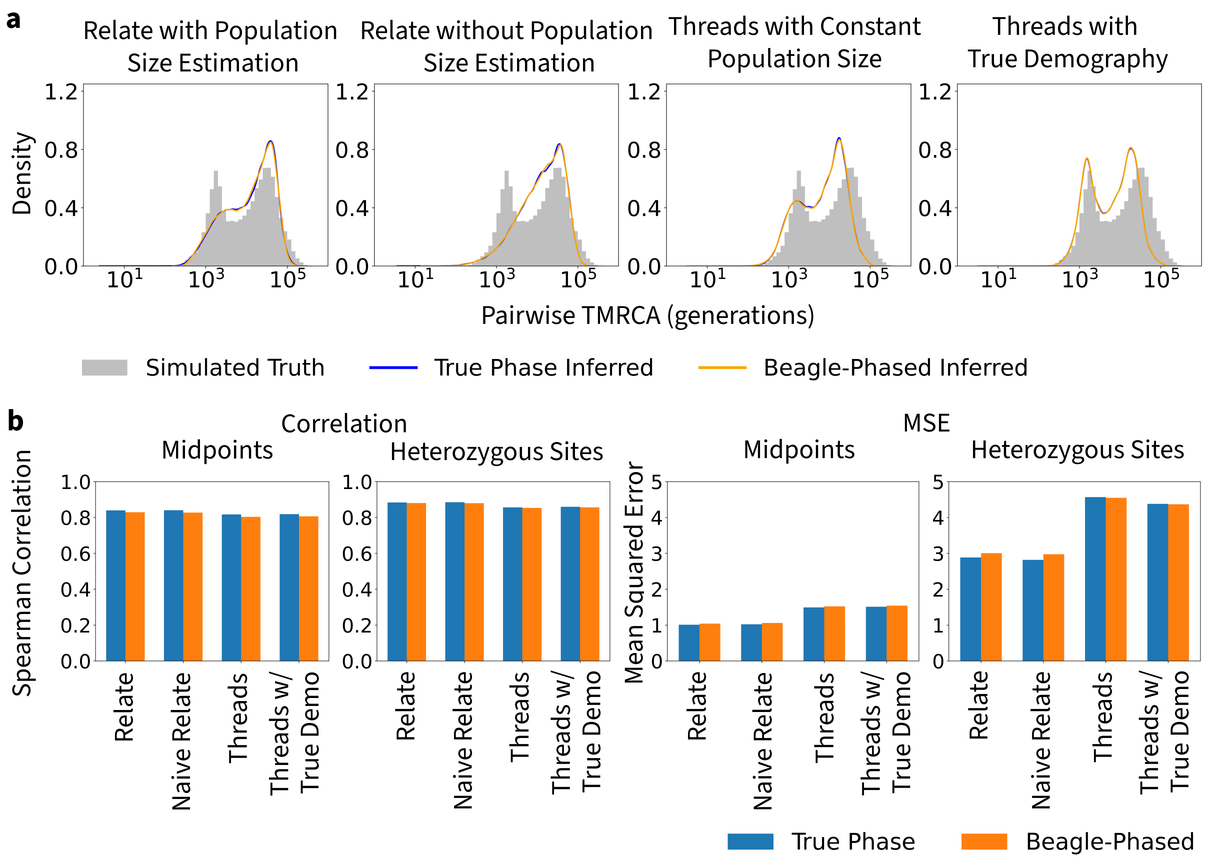


Figure 5: a. Distributions of pairwise coalescence times of true ARGs, ARGs inferred from true VCFs and ARGs inferred from Beagle-Phased VCFs by Relate with and without add-on population size estimation module and Threads with constant population size demography and true demography. b. The correlation (left half) and mean squared error (right half) between true ARGs and ARGs inferred by Relate with and without add-on population size estimation module and Threads with constant population size demography and true demography based on midpoint switch state reconstruction and at heterozygous sites.

### 3 Version Difference in Relate and tsinfer

In the main text, we presented the performance of the latest version tested for each method. However, we noticed that different versions of Relate and tsinfer produced different levels of recombination breakpoints, which would explain different relative performances observed in previous studies.

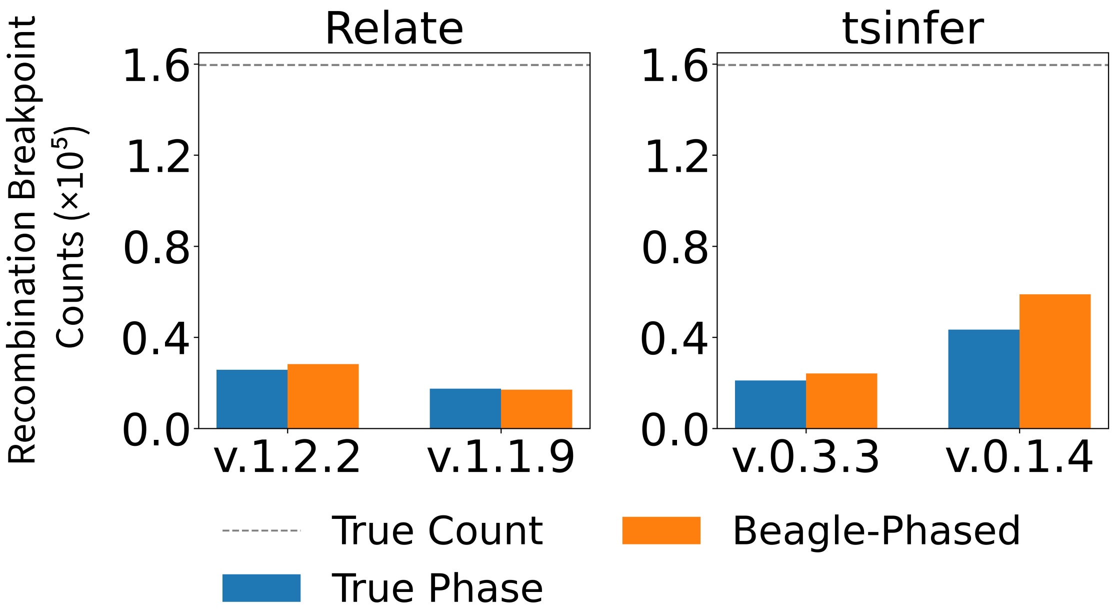


Figure 6: The counts of inferred recombination breakpoints in different versions of Relate (v.1.2.2 and v.1.1.9) and tsinfer (v.0.3.3 and v.0.1.4).

### 4 Sites under Selection in Anopheles

We presented one exemplary site under selection in the main text. The site was chosen arbitrarily among many sites in the cyp6p locus. Other nearby sites within the locus showed similar trends in inferred diversity and allele frequency trajectories as shown in Figure 7.

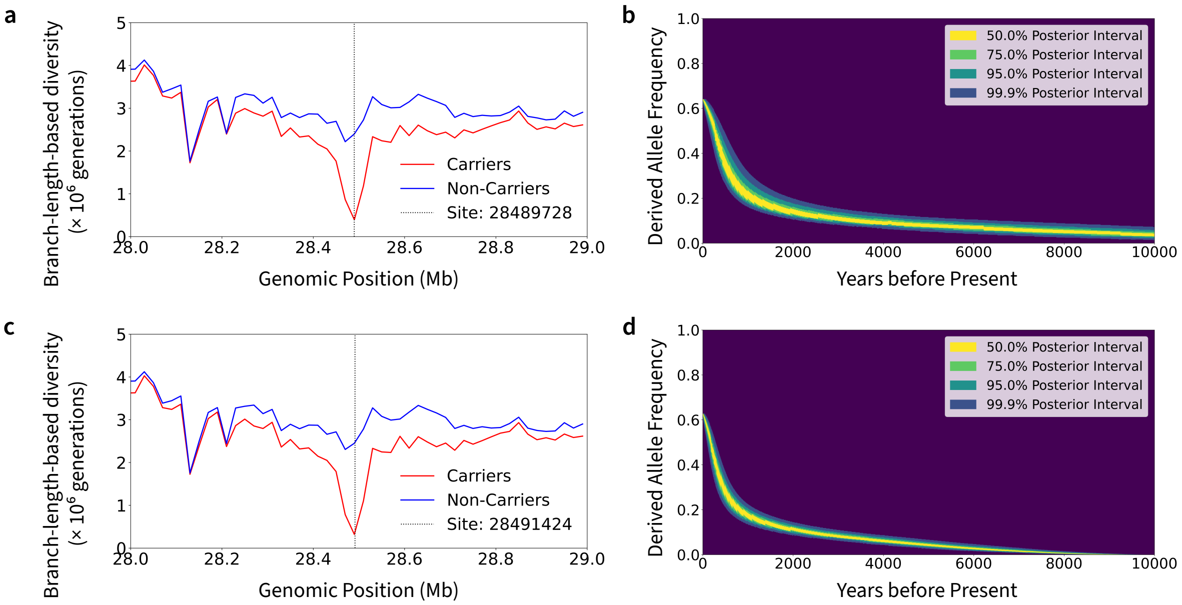


Figure 7: a. The branch-length-based diversity of carriers and non-carriers over the 1 Mb genomic region around the site (28489728) of selection in cyp6p. b. The CLUES2-estimated allele frequency trajectory of the mutation under selection (site 28489728). c. The branch-length-based diversity of carriers and non-carriers over the 1 Mb genomic region around the site (28491424) of selection in cyp6p. d. The CLUES2-estimated allele frequency trajectory of the mutation under selection (site 28491424).

### 5 Estimation of Mutation Age

We estimated the age of the focal mutation from 80 SINGER posterior samples. In each posterior sample, the focal mutation is placed on a branch bounded by two nodes. We assumed the mutation age has a uniform distribution between its bounds and averaged across the 80 posterior samples to obtain the posterior predictive distribution.

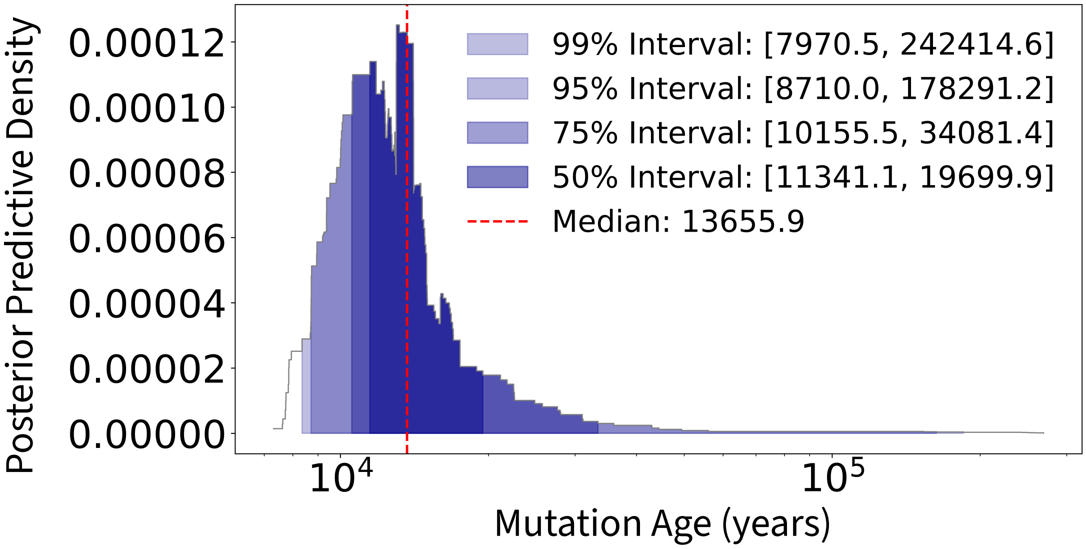


Figure 8: The posterior predictive density of the mutation age obtained from 80 SINGER posterior samples at site 28,487,111. The 50%, 75%, 95%, and 99% posterior predictive intervals (equal-tailed) are colored in the figure.
